# Isolation and Characterization of a Novel Calcium-Precipitating Alkali-Tolerant *Lysinibacillus sphaericus* in Local Urban Environment

**DOI:** 10.64898/2026.01.23.701334

**Authors:** Kah Wee Tan, Soon Keong Wee, Aden Yeoh, Zhenyu Zhang, Kang Hai Tan, Eric Peng Huat Yap

## Abstract

Concrete is an extensively used construction material in infrastructures due to cost-effectiveness and durability. However, its low tensile strength makes it prone to cracking, posing persistent maintenance challenges. Traditional repair methods are often expensive and difficult to implement, especially for critical structures. Recent studies propose a potential solution involving alkaliphilic or alkali-tolerant calcium-carbonate producing bacteria in aiding concrete crack repair through microbial-induced calcium carbonate precipitation (MICP) via urea hydrolysis. Considering the adaptive nature of bacteria, we hypothesize that naturally occurring concrete-inhabiting microbes possess properties crucial for mending cracks. Therefore, this study aims to isolate and evaluate these inherent microbes and their MICP ability. Six concrete samples were collected and cultured on Alkaline Nutrient Agar for microbial isolation. Urease activity was assessed phenotypically and genotypically. MICP activity was observed and validated under Scanning Electron Microscope/Energy Dispersive Spectroscopy (SEM/EDS). Out of the 49 isolates cultured, *Lysinibacillus sphaericus* PUMA0250 emerges as the most promising isolate for practical implementation, showing high pH survivability in an in-house formulated concrete agar medium (pH 12.6). *L. sphaericus* PUMA0250 exhibited MICP activity via urea hydrolysis and harbored all crucial genes involved in urease-mediated MICP. SEM/EDS analysis confirmed the structure and composition of the produced calcium carbonate precipitate. This study demonstrates the potential of *L. sphaericus* PUMA0250 to be incorporated into concrete to assess its healing efficiency. Its ability to thrive in the alkaline concrete environment and produce calcium by-products may aid concrete crack repair, opening microbiological avenues for sustainable and efficient concrete repair methodologies.

**IMPORTANCE:** Although concrete is widely used in construction, the construction material is prone to cracking, leading to costly repairs and structural degradation. This study isolated *Lysinibacillus sphaericus* PUMA0250 from tropical urban concrete, that can survive the highly alkaline environment typical of concrete structures in equatorial climate. Importantly, it produces calcium carbonate, a mineral that can help in concrete repair. These findings contribute to our understanding of how microbes that naturally produce calcium carbonate by-products can be used to repair concrete. For such applications, microbes must be able to withstand extreme conditions, including high pH and low nutrient availability. Our discovery of a resilient, native isolate highlights the potential of tropical concrete microbiomes as sources for self-healing construction technologies. This work lays the foundation for more sustainable, low-maintenance building materials and opens new possibilities in microbiology-driven solutions for urban infrastructure challenges.

## INTRODUCTION

Concrete, widely used in construction, faces significant challenges from crack formation and associated degradation caused by mechanical and environmental forces. These cracks compromise structural performance, serviceability and durability, while also accelerating steel corrosion (Sumra et al., 2020) due to ingress of moisture. Over time, factors like chloride ingress, alkali leaching, carbonation, corrosion, acid attacks, moisture and biodegradation can alter concrete’s pH, weakening its strength and durability. These changes increase the risk of structural failures, safety hazards and economic losses. Although timely repairs are crucial to prevent crack propagation, conventional methods like sealing, grouting and concrete replacement are often hindered by crack size, location and the operational demands of critical infrastructure.

Self-healing concrete using microbially-induced calcium carbonate precipitation (MICP) has drawn increasing interest since the discovery of calcium carbonate sedimentation by *Bacillus pasteurella* in the 1990s (Gollapudi et al., 1995). Among the MICP pathways, urea hydrolysis is widely studied due to its practical applications (Li et al., 2014, 2015). Ureolytic bacteria facilitate this process by breaking down urea in steps that ultimately precipitate calcium carbonate (CaCO_3_) (Zhao et al., 2022). This mechanism relies on urease-encoding operons (*ureABCEFGD*) for urease maturation and activity (Yap & Wong, 2019). Additionally, an effective MICP depends on components like proton-dependent ATP synthase (*atpABCDEFGH*), ammonium ion transport systems (*nrgAB*), urea transporters (*urtABCDE*), and nickel transporters (*hypAB*) (Mobley & Hausinger, 1989).

Concrete, despite its harsh environment characterized by high alkalinity, low nutrients, dryness and salinity, hosts a diverse microbiome. This includes *Proteobacteria*, *Firmicutes*, *Actinobacteria*, and other taxa such as *Cyanobacteria*, *Bacteroidetes*, *Acidobacteria*, and *Planctomycetes* (Kiledal et al., 2021). Fungi, including *Fusarium cerealis* and *Phoma herbarum*, have also been identified (Zhao et al., 2022). These microorganisms can be seeded from the materials used to cast concrete and adapt to environmental changes like calcite precipitation and seasonal shifts, causing the concrete microbiome composition to fluctuate (Kiledal et al., 2021). Their ability to modify their surroundings, either raising or lowering pH, influences their survival with both beneficial and detrimental effects (Ratzke & Gore, 2018). Concrete exhibits a highly alkaline environment with pH typically ranging from 12 to 13.8 (Sumra et al., 2020). Hence, considering the adaptive nature of bacteria, we hypothesize that naturally-occurring concrete-inhabiting microbes possess properties crucial for mending cracks. This study aims to isolate alkali-tolerant microbes found in concrete and characterize their MICP ability based on alkaline tolerance, ureolytic activity, and precipitate purity to assess their potential applications in self-healing concrete.

## RESULTS

### *Lysinibacillus sphaericus* PUMA0250 isolated from concrete samples

*Lysinibacillus sphaericus* PUMA0250, along with other 48 isolates, including *Bacillus infantis* PUMA0249, *Enterococcus casseliflavus* PUMA0251, *Paenibacillus lautus* PUMA0252, and *P. lautus* PUMA0253, were isolated from concrete samples obtained from buildings undergoing interior renovation (Table S1).

### *L. sphaericus* PUMA0250 is viable at pH 12.6

As *L. sphaericus* PUMA0250 was successfully cultured on alkaline nutrient agar (ANA) adjusted to pH 10 over a period of 72 hours at 30°C (Table S2), we characterized its viability in a highly alkaline environment. The growth curve study revealed that *L. sphaericus* PUMA0250 was viable in nutrient broth adjusted to pH 11 (Figure S1A). *L. sphaericus* PUMA025 had also demonstrated survival adaptation to a higher pH level characterized by a longer lag phase. However, no growth was observed in nutrient broth at pH 12, consistent with the challenges of a planktonic lifestyle under extreme alkalinity. To simulate *L. sphaericus* PUMA0250 viability on concrete, an in-house formulated concrete agar medium (CAM) of pH 12.6 was used as proxy to concrete. After 24 hours of incubation, microbial growth was identified as microcolonies (<1 mm) and was not visually distinct (Figure S1B). The microcolonies were viable after subculturing onto TSA from CAM, demonstrating the capacity to withstand the alkaline, oligotrophic conditions of a concrete proxy.

### *L. sphaericus* PUMA0250 precipitates calcium carbonate

The MICP capability of *L. sphaericus* PUMA0250 was evaluated by culturing the isolates in TSB supplemented with 2% urea for 4 hours to achieve log phase bacterial growth, followed by the addition of CaCl₂. Immediate formation of white precipitates was observed upon the introduction of CaCl₂. A notable increase in precipitate formation was evident in *L. sphaericus* PUMA0250 cultures supplemented with 2% urea (Figure S2). However, the dry weight (0.6390 g) of precipitates produced by *L. sphaericus* PUMA0250 did not significantly differ from those produced by two other isolates cultured from the concrete, namely, *L. fusiformis* T6-27-1 and *L. fusiformis* S6-15-1 (Table 1). Given this similarity, further analyses on crystal morphology and elemental composition were necessary to determine whether the precipitates were the result of urease-mediated calcium carbonate formation. Using scanning electron microscopy (SEM) coupled with energy-dispersive X-ray spectroscopy (EDS), the precipitates from *L. sphaericus* PUMA0250 and *L. fusiformis* T6-27-1 exhibited colocalization of calcium, carbon, and oxygen, consistent with CaCO₃ formation (Figure 1). In contrast, the precipitates from *L. fusiformis* S6-15-1 showed colocalization of calcium and chloride, suggesting the presence of calcium chloride likely from the media (Figure 1). EDS analysis confirmed the elemental composition of the precipitates produced by *L. sphaericus* PUMA0250, revealing atomic percentages of 7.8% calcium, 35.4% carbon and 39.1% oxygen, strongly indicating that the precipitates were predominantly calcium carbonate (Table 1).

**Table 1.**
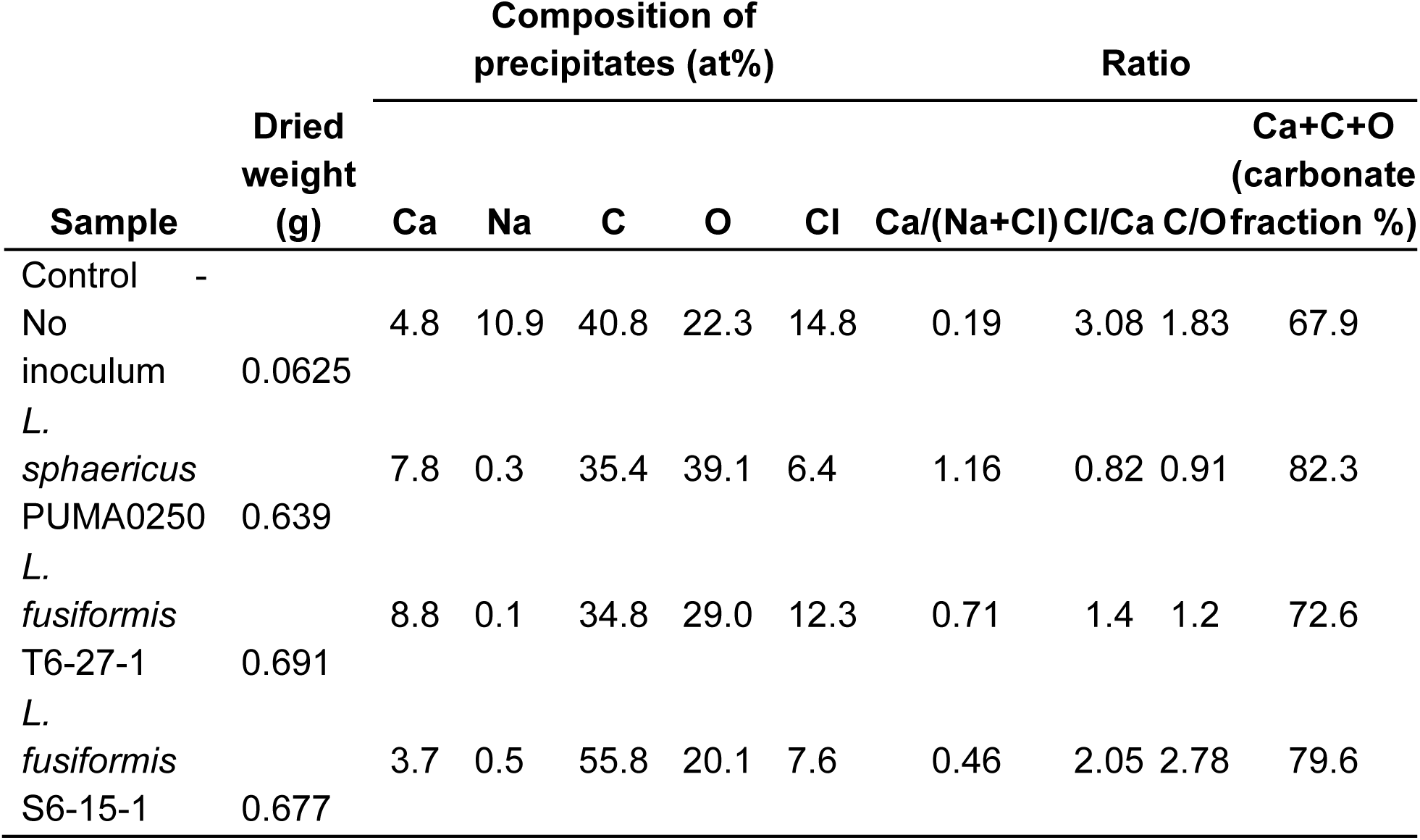
EDS Analysis. Atomic percentage of elemental composition of dried precipitate produced by various isolates (*L. sphaericus* PUMA0250, *L. fusiformis* T6-27-1 and *L. fusiformis* S6-15-1) after 2% urea supplementation.

**Figure 1.**
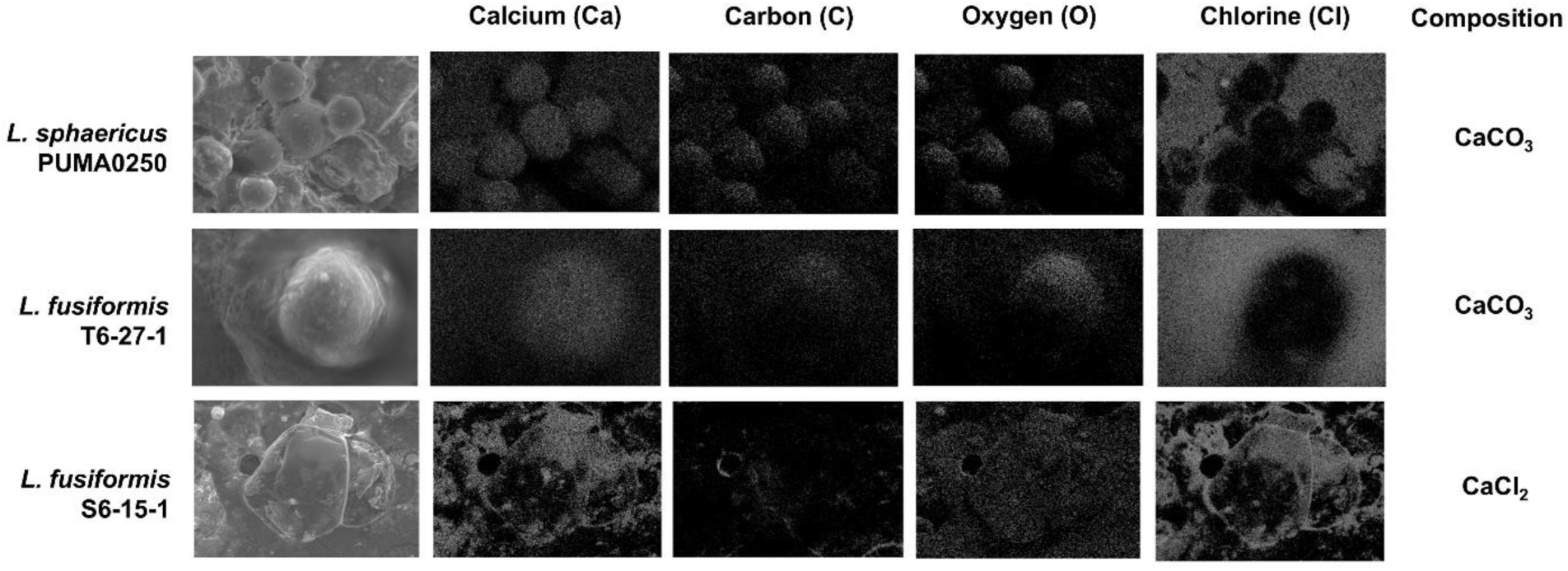
SEM-EDS analysis of precipitates produced by bacterial isolates after 2% urea supplementation.

### *L. sphaericus* PUMA0250 undergoes MICP via urea hydrolysis pathway

As calcium carbonate precipitation can be achieved through urea hydrolysis (Brink, 2010), it is crucial that microbes possess urease-producing characteristics for MICP to occur. The isolate *L. sphaericus* PUMA0250 was identified to possess urease-producing characteristics as the color of the Christensen’s urea agar turned from yellow to fuschia (Figure S3). *L. sphaericus* PUMA0250 has exhibited to be a rapid urease-producing isolate as the phenotypic results were concordant with the control using *P. mirabilis* C0472, which is known to have a rapid urease-producing activity (Brink, 2010). This suggests that *L. sphaericus* PUMA0250 has efficient MICP capability through urea hydrolysis pathway. Conversely, *L. fusiformis* T6-27-1 and *L. fusiformis* S6-15-1 have delayed urease producing activity that were similar to *Klebsiella pneumoniae* C0290 control isolate (Brink, 2010).

### *L. sphaericus* PUMA0250 harbored all crucial urease genes (*ureABCEFGD*)

*L. sphaericus* PUMA0250 has a genome size of 4,552,253 bp with GC content of 37%. The draft genome has a total of 75 contigs with N50 of 239.7 kb. *L. sphaericus* PUMA0250 had a dDDH_4_ similarity of 96% to *L. sphaericus* CBAM5 and similarity of 99.4% and 99.6% using the ANIb and ANIm method respectively (Table S2). However, it has a dDDH_4_ similarity of 23.6% to *L. sphaericus* KCTC 3346, below the 70% threshold used for species demarcation, indicating that PUMA0250 may represent a novel species or a highly divergent lineage. The publicly available genomes of *L. sphaericus* were examined and found to have 2 broad subspecies (Figure S4).

*L. sphaericus* PUMA0250 possesses a urease operon containing structural genes *ureA, ureB* and *ureC,* as well as the accessory genes *ureE, ureF, ureG* and *ureD,* which facilitate urease maturation (Yap and Wong, 2019). Comparative analysis with other reported species showed that the orientation of the urease genes in *L. sphaericus* PUMA0250 has high conservation to that of the *L. sphaericus* OT4b.25 and *Sporosarcina pasteurii* BNCC 337394 strains (Figure 2). Given the conserved synteny of the urease operon among phylogenetically related strains known to exhibit ureolytic activity, supports the likelihood that the operon in *L. sphaericus* PUMA0250 is functional.

**Figure 2.**
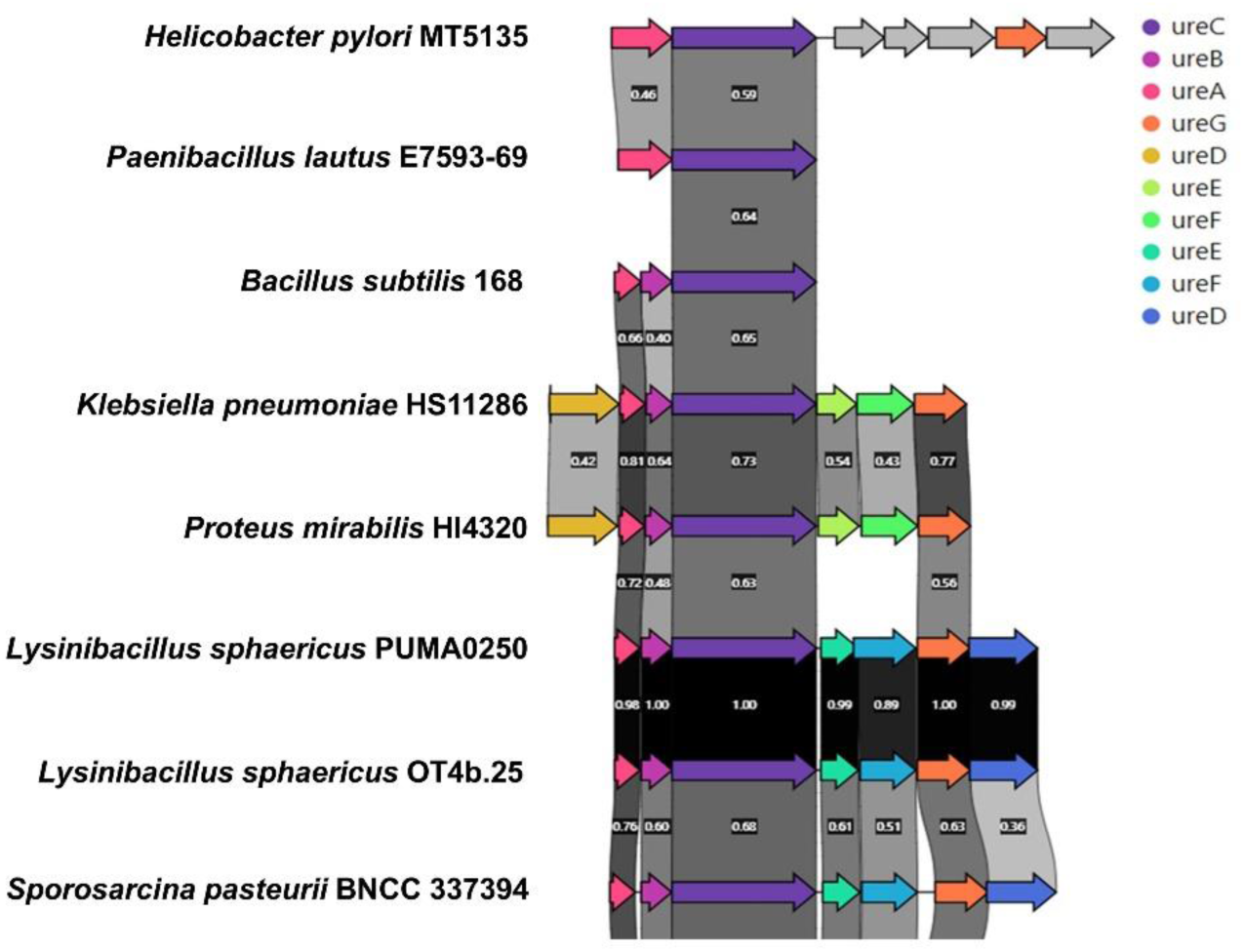
Genomic orientation of urease operon (*ureABCEFGD*) in *L. sphaericus* PUMA0250 visualized using Clinker (Gilchrist & Chooi, 2021).

### Presence of other essential genes for MICP in *L. sphaericus* PUMA0250

The possession of urease genes alone is insufficient for MICP to occur. Additional components like proton-dependent ATP synthase, ammonium ion transport systems, energy-dependent urea transporters, and energy-dependent nickel transporters are also important for MICP to occur. *L. sphaericus* PUMA0250 isolate likely undergoes a series of intracellular pathways for MICP to be successfully carried out (Figure 3). It was found to have all the essential proton-dependent ATP synthase genes (*atpABCDEFGH*) and ammonium ion transport systems (*nrgAB*). Conversely, it lacks the energy-dependent urea transporter operon (*urtABCDE*), which is crucial for the active transport of urea into the cells (Figure 3). This suggests that urea transport in *L. sphaericus* PUMA0250 may rely on passive diffusion (Veaudor et al., 2019). *L. sphaericus* PUMA0250 possesses a high-affinity nickel transporter operon (*nikABCDE*) instead of the energy-dependent nickel transporter operon (*hypAB*) to facilitate the intracellular transport of nickel (Figure 3). Together, these findings suggest that *L. sphaericus* PUMA0250 has the genetic potential to perform ureolytic MICP.

**Figure 3.**
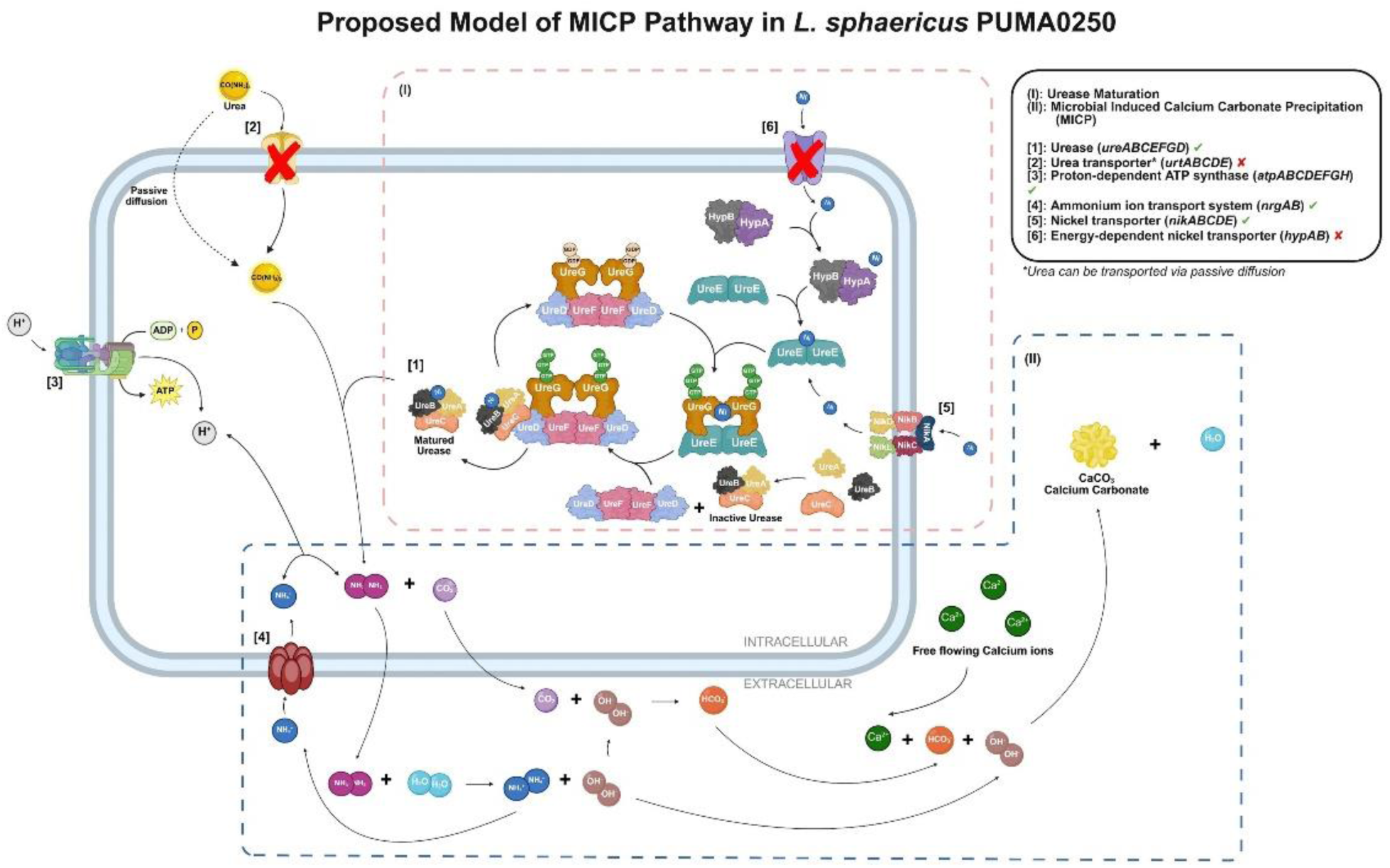
Schematic diagram of the urease maturation pathway (Yap & Wong, 2019) and microbially-induced calcium carbonate precipitation (MICP) process through urea hydrolysis and urea transportation (Mobley & Hausinger, 1989) in *L. sphaericus* PUMA0250 created with BioRender.com. The genes that were not detected in *L. sphaericus* PUMA0250 are indicated with a red cross.

## DISCUSSION

### Alkali-tolerant bacteria are found in concrete

Concrete is a widely utilized material in the construction industry, and during its hydration process, the pH can increase significantly, reaching up to pH 13.8 (Natkunarajah et al., 2022). Such an extreme alkaline environment is generally uninhabitable for most microbes. Our study is limited by employing a convenient random sampling approach and constrained by the aerobic growth culture conditions, which would exclude anaerobic microbial species with self-healing potential in these harsh environments. However, microbes were still able to be isolated with an extended incubation period of up to 72 hrs, suggesting that alkali-tolerant microbes require additional time to adapt to the high pH conditions before they could thrive. This is illustrated when *L. sphaericus* PUMA0250 had a longer lag phase when cultured under higher pH conditions (Figure S1A). This could involve the expression of acidic compounds to maintain the cytoplasmic pH at levels lower than the external pH (Saito & Kobayashi, 2003). Alternatively, the presence of different populations of *L. sphaericus* with varying pH tolerance levels might explain the observed phenomenon. There may be a smaller subset of *L. sphaericus* that exhibits a higher pH tolerance (pH 11) as compared to the overall populace in the sample, necessitating a longer time for these microbes to thrive due to competition for growth with a larger population of lower pH tolerance. We observed that most of the isolated microbes were identified as belonging to the *Bacillus* genus (Table S2) which is commonly known to have an innate ability to produce endospores. Given that *L. sphaericus* is phylogenetically closely related to this group, it is highly probable that it also employs endospore formation as a key survival mechanism. This viability is directly demonstrated by its ability to form microcolonies on an in-house formulated concrete agar medium (CAM) at pH 12.6, and crucially, to remain viable after subculturing from this harsh environment back to TSA. These endospores serve as protective structures for bacterial cells to survive in harsh environments, including extreme pH gradients and limited nutrient availability like concrete (Basta & Annamaraju, 2024). Therefore, the extended lag phase observed in the growth curve study likely represents not just physiological adaptation, but also the temporal process of endospore germination and the outgrowth of viable cells once conditions become more favorable.

### *PUMA0250* is a member of Lysinibacillus sphaericus taxonomic group 1

In this study, it was found that *L. sphaericus* have 2 broad subspecies (Figure S4). Based on various comparison methods (dDDH_4_, ANIb, and ANIm), PUMA0250 is closely related to *L. sphaericus* CBAM5, an entomopathogenic strain under taxonomic group 1 (Peña-Montenegro et al, 2015). However, it is widely acknowledged that a limited number of sequenced genomes might not accurately reflect the entire genetic repertoire of a species (Deng et al., 2010; Medini et al., 2005), prompting a suggestion to re-evaluation *L. sphaericus* as a species (Gómez-Garzón et al., 2016). Early studies relying on less sensitive methods have hindered the estimation of diversity and created heterogenous species that include toxic and non-toxic strains (Nakamura, 2000). As such, the variability in toxicity might arise due to genetic variability and misclassification (Gómez-Garzón et al., 2016). *L. sphaericus* OT4b.49 was characterized through the mosquito larvicidal toxin genes it possessed (Gómez-Garzón et al., 2016). Although WGS revealed the absence of *binA, binB, cry48*, and *cry49* genes in *L. sphaericus* PUMA0250, *mtx2* toxin-encoding genes were present. However, further analysis is needed to confirm its entomopathogenic nature. Furthermore, as *L. sphaericus* PUMA0250 was inherently inhabiting concrete, exploring its bioremediation potential through pan-genome analysis could be a potential way to describe this species.

### *L. sphaericus* PUMA0250 is a potential candidate for MICP-based bioconcrete

The mitigation of cracks in concrete structures is currently accomplished through conventional solutions, such as crack injection, concrete patching and resurfacing, reinforcement installation, crack mitigation and slab jacking. However, these methods are plagued by issues of high cost, time-consuming implementation, and lack of sustainability (Palmer, 2009; Roels et al., 2022). The persistent growth of cracks within concrete structures leads to spalling and the corrosion of reinforcing bars, further accelerating structural deterioration (Mancio et al., 2004; Tannous, 1997; Virmani & Clemena, 1998). Therefore, these issues often result in significant replacement expenses and contribute to environmental challenges (Nkurunziza et al., 2005; Van Tittelboom et al., 2010; Xu et al., 2015). Addressing these limitations calls for the exploration and development of alternative crack management strategies in concrete that are both cost-effective and environmentally conscious, meeting the demands of modern construction practices. As such, incorporating suitable bacteria into concrete could be a more effective and sustainable approach.

*L. sphaericus* PUMA0250 is a potential candidate for self-healing concrete due to its high pH tolerance (pH 12.6) and MICP activity. The calcium carbonate precipitates produced by *L. sphaericus* PUMA0250 are byproducts essential for the mending of cracks in concrete. However, its self-healing capabilities directly in a harsh concrete environment and its healing efficiency must be further assessed. This study henceforth opens numerous downstream development opportunities, such as the potential of incorporating *L. sphaericus* PUMA0250 into concrete mixtures through microcapsule technology (Han et al., 2021) or the application of a microbial spray on concrete surfaces with existing cracks. These innovative approaches hold promises for enhancing the self-healing capabilities of concrete structures, contributing to increased durability and sustainability in the construction industry. A culturomics approach may be pursued to optimize and standardize the concrete agar medium for the growth and activity of *L. sphaericus* PUMA0250, ensuring its effectiveness in self-healing concrete applications. Besides, the metabolic conversion of calcite by ureolytic microbes would result in an increase in pH of the environment (Zhao et al., 2022) which could be disadvantageous to the viability of microbes. Hence, downstream studies can be carried out by performing calcium carbonate precipitation in a high pH environment to assess its implications on microbial growth.

In this study, we have demonstrated the presence of ureolytic bacteria in concrete indicating their adaptation to the harsh environment. One such novel strain, *L. sphaericus* PUMA0250, has been shown to have potential applications for mending concrete cracks given its ability to thrive in high alkaline environment with pH up to 12.6 and urease-producing activity on Christensen’s urea agar. It harbored all the essential genes involved in MICP and could produce CaCO_3_ by-product through urea hydrolysis. There are numerous promising downstream development opportunities for microbiologically mediated self-healing concrete. These future research directions will help to refine and validate the practical application of *L. sphaericus* PUMA0250 in construction, contributing to the development of innovative and sustainable self-healing concrete technologies.

## MATERIALS AND METHODS

### Field sampling of concrete

A total of six concrete samples were taken from various interior areas of two renovating residential apartments in Singapore. At least 20g of these samples were carefully obtained and placed into a 50mL centrifuge tube. Tubes were labelled and details (photos, source, type of source, type of apartment, location, elevation, interior location & description) of the samples were recorded.

### Enrichment of microbes found in concrete samples

A total of 15g of each concrete sample was washed and vortexed with 15mL of phosphate-buffered saline with 0.1% Tween-20 (PBS-T) before setting aside to allow concrete sediments to settle. Next, 5mL of supernatant from each sample was enriched with 5mL of Luria-Bertani (LB) broth and incubated overnight in a shaker at 37°C at 200rpm.

### Isolation of alkaliphile and alkali-tolerant microbes

Culture isolation of alkaliphilic microbes was performed using spot plating technique. The enriched samples were serially diluted in PBS by 3- to 5- log orders of magnitude (ie. 1,000X to 100,000X fold dilution). Spot plating was done using 20μl spots onto Alkaline Nutrient Agar with different pH values (pH 9, 10, 11 & 12) and Malt Extract Agar (pH 4.92, 9, 10, 11 & 12). The culture plates were incubated at 30°C for 24 hours and subsequently till 72 hours for slow growing microbes. Isolates with different morphology were sub-cultured twice onto Tryptic Soy Agar (TSA) with incubation at 30°C for 24 hours to obtain axenic cultures.

### Bacterial Identification via 16S ribotyping

The 16S rRNA gene was amplified using colony quantitative polymerase chain reaction (qPCR). Isolates were chosen with priority for those that grew in higher pH conditions (pH 11 & 12), followed by their morphological differences. Colony PCR templates were prepared by emulsifying a single colony into 50μl of molecular grade water and heat-lysed at 95°C for 10 minutes. qPCR was conducted by QuantStudio 6 Flex (Applied Biosystems, Life Technologies) in a 50μl reaction mix containing 0.14μM of each primer (27F: 5’-AGAGTTTGATCMTGGCTCAG-3’ and 1492R: 5’-TACGGYTACCTTGTTACGACTT-3’) (Frank et al., 2008), 1X KAPA2G Fast HotStart ReadyMix (KAPA Biosystems, USA), 1X EvaGreen® Dye (Biotium, USA). The cycling conditions were initial denaturation at 95°C for 3 min, followed by 40 cycles of 95°C for 15s, 60°C for 15s and 72°C for 5s, and a final extension step of 72°C for 1min. PCR amplicons were purified using Agencourt AMPure XP PCR Purification kit (Beckman Coulter) following manufacturer’s instructions. The purified amplicons were subsequently sent out to 1st BASE (Singapore) for single pass DNA sanger sequencing using the universal primers 27F and 1492R.

Both forward and reverse reads were analyzed and curated manually based on their Phred quality score to obtain the consensus sequence using SeqTrace 0.9.0. The consensus sequence was uploaded onto NCBI (National Center for Biotechnology Information) Basic Local Alignment Search Tool (BLAST) to identify the bacteria species against the 16S ribosomal (rRNA) sequences database.

### Screening for urease-producing alkali-tolerant isolates

Urease production was assessed using Christensen’s urea agar (Brink, 2010), prepared by aseptically adding urea base (1g/L peptone, 1g/L dextrose, 5g/L NaCl, 2g/L KH_2_PO_4_, 20g/L urea, 0.012g/L phenol red, 100mL of milli-Q water) to 900mL of autoclaved BactoTM agar (BD) solution through a 0.45mm pore size filter. 5mL of Christensen’s urea agar solution was distributed into each 15mL falcon tube and allowed it to solidify as a slant agar. Previously cryostored isolates were subcultured onto TSA and incubated at 37°C overnight. A single colony of each isolate was picked and streaked onto the Christensen’s urea agar and was incubated in 37°C overnight. Rapid urease producing *Proteus mirabilis* C0472 and delayed urease producing *Klebsiella pnuemoniae* C0290 were used as positive controls while *Escherichia coli* K12 and no inoculum were used as negative controls. The control strains were obtained from LKCMedicine Microbial Repository.

### Determination of calcium carbonate precipitation activity

A total volume of 0.63mL of urease-producing bacterial culture with optical density (OD) reading of 0.2 value at 600nm was inoculated into 13mL of TSB supplemented with 2% of urea and incubated at 37°C in a shaker at 200rpm for 4 hours. A volume of 10mL of the culture broth, with and without inoculum was used for the determination of the CaCO_3_ precipitation activity. *E. coli* K12 was chosen as a negative control for its known non-urease producing properties. The bacterial cells were removed by centrifuging the culture at 9000rpm for 10min. CaCl_2_ solution was subsequently added to the supernatant to a final concentration of 0.4M CaCl_2_. The resulting CaCO_3_ precipitates were collected, washed with deionized water, and dried at 90°C until a constant weight was attained for measurement.

### Scanning Electron Microscopy – Energy Dispersive Spectroscopy (SEM-EDS)

The dried CaCO_3_ precipitate and its elemental composition were assessed by SEM-EDS using JSM-7600F Schottky Field Emission Scanning Electron Microscope (JEOL Ltd., Singapore) with 30kV accelerating voltage.

### Assessing pH tolerance of urease-producing alkali-tolerant isolates

Each isolate was cultured by inoculating a single colony into 25mL of nutrient broth (pH 7) and incubated overnight at 37°C in a shaker at 200rpm. 15μl of overnight culture was added to 25mL of fresh nutrient broth (pH 7) and was incubated at 37°C in a shaker at 200rpm for 2 hours. Each fresh culture was diluted 3X with nutrient broth of various pH (pH 9, 10, 11 & 12) in a 96-well plate to achieve 10^5^ cells/mL. The growth kinetics of the isolates were performed in triplicates and were observed using a Synergy H1 microplate reader (Biotek) with continuous shaking at 37°C. The absorbance at a wavelength of 600 nm was measured to determine the optical density at 10min interval for 24 hours. The growth curves of each isolate in varying pH conditions were plotted using GraphPad Prism 5.

### Microbial viability on in-house formulated concrete agar medium

Cement base mixture, containing 100g/L of sand (top sand from a local park in Singapore), 200g/L of cement (NS Portland Composite Cement), and nutrient agar (13g/L nutrient broth powder and 13g/L of agar powder) was autoclaved and aseptically mixed to create an in-house formulated concrete agar medium (CAM) in the ratio of in the ratio of 1:2:7. Concrete agar medium was equally distributed to each petri dish with intermittent shaking to ensure homogenous mixture. Urease producing microbes were inoculated onto the in-house formulate concrete agar medium and was incubated at 37°C overnight. Colony from the concrete agar medium was streaked onto TSA and incubated at 37°C overnight for growth verification.

### Genomic characterization of urease-producing alkali-tolerant microbes

Bacterial isolates for whole genome sequencing were incubated on TSA from the cryostock at 37°C overnight. Colonies were picked and DNA extraction was performed using QIAamp® DNA Mini Kit (Qiagen). Quality and concentration assessment of the extracted DNA was performed using Nanodrop 2000c Spectrophotometer (Thermo Fisher Scientific, USA) and Qubit HS dsDNA Assay Kit on Qubit 4 Fluorometer (Thermo Fisher Scientific, USA). Whole genome sequencing was done on Illumina NovaSeq using 2×151bp paired-end configuration.

*De novo* assembly was performed with Unicycler Galaxy version 0.5.0 (Wee et al., 2023; Wick et al., 2017; Wee & Yap, 2021). Species determination was done by dDDH using TYGS server (Meier-Kolthoff & Göker, 2019). Genes were annotated using NCBI Prokaryotic Genome Annotation Pipeline 4.9 (Tatusova et al., 2016) and gene synteny was analyzed using Clinker (Gilchrist & Chooi, 2021). Default parameters were used for all software tools.

The draft genome assemblies of the isolates, *L. sphaericus* PUMA0250 together with *B. infantis* PUMA0249, *E. casseliflavus* PUMA0251, *P. lautus* PUMA0252 and *P. lautus* PUMA0253, were submitted to the GenBank database under accession number GCA_036620755.1, GCA_036620775.1, GCA_036620735.1, GCA_036620705.1 and GCA_036620695.1 respectively.

## ACKNOWLEDGMENTS

This research was funded by the Singapore Ministry of Education Academic Research Fund RG87/21, Research Centre for Excellence IDMxS, and Covid-19 grant from Lee Kong Chian School of Medicine. K.W.T. was supported by a Nanyang Technological University – Ma Kuang Scholarship.

